# Dynamics in the assembly of the 30S ribosomal subunit investigated by coarse-grained simulations

**DOI:** 10.1101/2023.06.13.544708

**Authors:** Xin Liu, Zhiyong Zhang

## Abstract

The ribosome is a large biomolecular complex responsible for protein synthesis. In *Escherichia coli* (*E. coli*), a complete ribosome is composed of a 30S small subunit and a 50S large subunit. For about half a century, the 30S subunit has been a key model system for studying the *in vitro* assembly of the ribosome, and an assembly map has been proposed. However, structural details in the assembly of this protein-RNA complex remain elusive. In this paper, we have conducted a series of coarse-grained simulations following the order of the assembly map, in order to investigate conformational dynamics during the assembly process of the 30S subunit. It has been found that, the tertiary structure of the naked 16S rRNA is very unstable, and that is the case after binding of the early-assembly proteins. The mid-assembly proteins can significantly restrict the mobility of the 16S rRNA and make the latter close to the native structure. The final binding of the late-assembly proteins would fully obtain the collective motion of the 16S rRNA. In particular, proteins S9 and S3 may have more important contributions to the assembly of the 30S subunit than other S proteins. Our strategy of coarse-grained simulations can be generally used to study assembly dynamics of large biomolecular complexes as long as the assembly map is available.

## Introduction

The ribosome is a large biomolecular complex including proteins and RNAs, which is responsible for catalyzing protein synthesis^[1]^. A complete ribosome contains a large subunit and a small subunit. In prokaryotes and archaea, the 50S large subunit is composed of a 23S rRNA, a 5S rRNA, and 33 proteins, whereas the 30S small subunit consists of a 16S rRNA and 21 proteins^[2]^.

The evolution of a large biomolecular complex can be seen as a long period of assembly, and so the assembly can reflect the pathway of complex evolution^[3]^. The assembly process of a complex is dynamic^[4]^, which is related to the order in which the same or different components come together^[3] [5, 6]^. The assembly of the ribosome in vivo requires hundreds of assembly factors to work together under certain conditions, but the assembly in vitro can be done without any assembly factors^[7]^, which has been studied for a long time. As early as 1960s, the in vitro spontaneous assembly ability of the 30S subunit in the *E. coli* ribosome was confirmed by the Nomura laboratory^[8]^, and subsequently, the assembly of the 50S subunit was confirmed by the Nierhaus laboratory^[9]^. Later, the ability of the ribosome from other bacteria to assemble into catalytically active structures through separate natural RNA and proteins has also been demonstrated^[10-15]^.

The 16S rRNA in the 30S ribosomal subunit has four independent domains: the 5’domain, the central domain, the 3 ’ major domain, and the 3 ’ minor domain. Each of the first three main domains can be independently assembled with the corresponding S proteins in vitro^[16-18]^. The naked 16S rRNA is very active and requires the assembly of the S proteins to form a stable ordered structure. The 30S subunit can be reconstituted in vitro with the 16S rRNA as a starting point by adding the necessary S proteins, so that exhibit their relevant biochemical characteristics^[19]^. Under reconstitution conditions, several methods of adding proteins to the 16S rRNA using combined binding allow the S proteins to be divided into three classes to form an assembly map^[20]^. Primary binding proteins bind directly and independently to the 16S rRNA, which are thought to initiate the folding of each of the three main domains. Secondary binding proteins require at least one primary binding protein before binding to the 16S rRNA, and tertiary binding proteins require at least one protein from the first two stages to be assembled. Through in vitro assembly kinetic experiments, a kinetic assembly map was formed, and the S proteins were divided into early-, mid-, mid-late- and late-assembly proteins. The kinetic assembly data reflect various aspects of domain assembly and binding trends, indicating that the assembly ranges from the 5′ domain to the 3′ domain, which are consistent with the co-transcriptional assembly^[21]^.

In recent years, various experimental techniques have been used to study the assembly of the 30S subunit. Electrospray ionization mass spectrometry (MS), along with the latest cryo-EM techniques, can visualize intermediates at near-atomic resolution throughout the pathway of subunit construction and is therefore widely used to study the assembly pathway of complexes^[5]^. While such data has proven to be a very useful starting point, it does not provide detailed information about dynamics in the assembly process. That is, only the structure can be obtained, and there is not enough information of how the complex form the existing structure.

Molecular dynamics (MD) simulations have long been used in studying dynamic processes of large biomolecules. They are generally atomic based on classical mechanics as kinetic principles, and molecular mechanical force fields are constantly improving^[22]^. However, all-atom MD is very expensive in simulating large biomolecular complexes such as the ribosome, and it would be difficult to describe the complete and long time-scale kinetic process^[23]^. In recent years, coarse-grained (CG) models have been used to reveal the physical mechanism of a series of protein/nucleic acid molecules^[24-28]^ because their computational efficiency is significantly higher than all-atom MD. CG simulations have been continuously optimizing and calibrating relevant force fields and methods due to their ability to display processes such as assembly of large biomolecular complexes at long time-scale^[29]^. There are many CG model available, such as the structure-based model^[30]^, the multi-scale coarse-grained (MS-CG) method^[31]^, the MARTINI model^[32, 33]^, and the SAFT model^[34]^.

Therefore, in this work, based on the kinetic assembly map, we study structural dynamics in the assembly process of the 30S subunit through CG simulations. The simulation data is compared with the existing experimental data of in vitro assembly, which can reveal the role of the specific S proteins in the assembly process.

## Materials and methods

### Simulated systems

The 30S ribosomal subunit of *E. coli* is composed of a 16S rRNA and 21 ribosomal proteins (S proteins, named S1∼S21 in a decreasing order of molecular weight). An atomic model of the 30S subunit was taken from^[35]^, in which the 16S rRNA contains 1536 nucleotides and 20 S proteins (S2-S21) are available (**Fig. 1**). The three main domains (5’, central, and 3’ major) of the 16S rRNA make up the body, the platform, and the head, respectively. The assembly of the S proteins are divided into four stages according to the kinetic map (**Table 1**). The early-assembly proteins include S4, S6, S11, S15, S16, S17, S18, and S20 (**Fig. 1**, magenta), the mid-assembly proteins include S7, S8, S9, S13, and S19 (**Fig. 1**, orange), the mid-late-assembly proteins include S5 and S12 (**Fig. 1**, blue), and the late-assembly proteins include S2, S3, S10, S14, and S21 (**Fig. 1**, red).

**Table 1.** Kinetic assembly map of the S proteins in the 30S ribosomal subunit

| Group | Protein |
| --- | --- |
| **I (early)** | S4 |
|  | S6 |
|  | S11 |
|  | S15 |
|  | S16 |
|  | S17 |
|  | S18 |
|  | S20 |
| **II (mid)** | S7 |
|  | S8 |
|  | S9 |
|  | S13 |
|  | S19 |
| **II' (mid-late)** | S5 |
|  | S12 |
| **III (late)** | S2 |
|  | S3 |
|  | S10 |
|  | S14 |
|  | S21 |

**Fig. 1.**
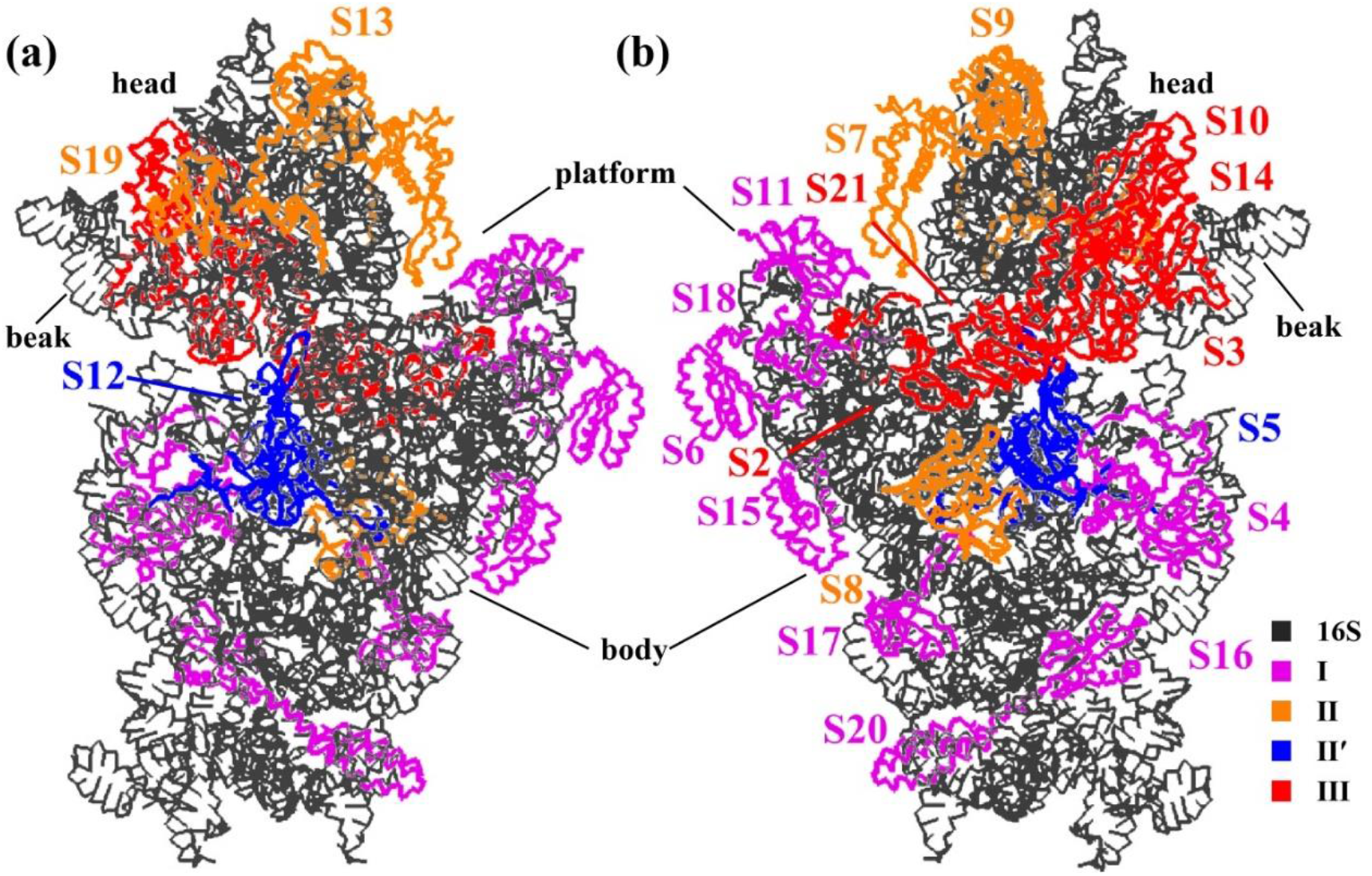
Structure of the 30S ribosomal subunit. (a) The front side (the interface with the 50S subunit), and (b) the back. The 16S rRNA is colored in black, and the S proteins are colored differently according to the kinetic assembly map (Table 1). The body, platform, head, and beak are labeled.

We built the following systems to run CG simulations. (1) The naked 16S rRNA that is the starting state of the assembly. (2) The 30S subunit that is the end state of the assembly. (3) Between the starting and the end states, we added the S proteins one by one, following the order shown in Table 1.

### The CG models

In the off-lattice Gō model of a protein, each amino-acid residue is represented by a CG particle, most often at the position of its C_α_ atom. The potential energy function is:

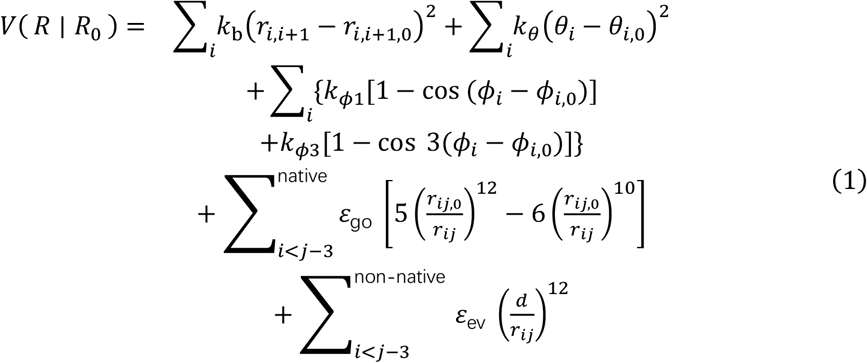

The first three terms represent potentials for virtual bond lengths, bond angles, and dihedral angles, respectively. The fourth term is the native contact potential between non-local residue pairs, where the summation is restricted to those pairs in the reference structure. The last term represents an excluded volume effect that is a penalty for those non-native contacts.

In this work, a Gō-like protein model called AICG2+^[36]^ was used:

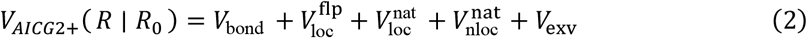

V_*bond*_ is the first term of Eqn. 1. 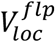 is a generic flexible local potential for the virtual bond angles and dihedral angles. 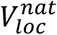 is the structure-based local potential. 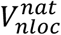 is the structure-based nonlocal contact potential, which is similar to the fourth term of Eqn. 1 except that these ε_go_ are not uniform but determined from atomic interactions between specific residue pairs. V_*exv*_ is the last term of Eqn. 1. Additionally, electrostatic forces were computed using the Debye–Hückel equation with a cut-off of 20 Å and ion strength of 0.15 M. The dielectric constant was set as 78.0 F/m. The off-lattice Gō model^[37]^ and its derivatives represent quasiharmonic fluctuations near native structures while achieving a perfect funnel energy landscape.

In the 16S rRNA, each nucleotide is represented by three CG particles, which are phosphate (P), sugar (S), and base (B), respectively. This model is a rational treatment to chemical distribution of nucleotide. The potential is described by the following terms: the local potential, the non-local contact interaction, the excluded volume term, and the electrostatic interaction. Since the 16S rRNA is largely double-stranded, a PAIR_RNA potential needs to be included to maintain the stability of base pairs. This interaction has a similar form as the general contact interaction, but use different coefficients discriminated by number of base-paring hydrogen bonds.

For interactions between the 16S rRNA and the S proteins, we also used the G ō model with the addition of repulsion and electrostatic interactions.

### CG simulations

All the CG simulations were conducted by CafeMol version 3.2.1^[30]^ using Langevin dynamics, with a time step of 0.3 in CafeMol time units. Each CG simulation was run for 10^8^ steps, and the temperature was set to 300 K. The overall translation and rotation of the entire molecule were allowed.

go_unit is an overall scaling factor of the pairwise interaction strength. For each native contact (intra-molecule or inter-molecule), the default value of go_unit is 1.0. We need to determine a proper value of go_unit, in order to observe significant conformational dynamics of the 30S subunit, while preserve its tertiary structure. Therefore, CG simulations of the 30S subunit were conducted by trying different values of go_unit, and three independent simulations were run for each go_unit. It has been found that a go_unit of 0.8 is appropriate. We then fixed the go_unit as 0.8 for both intra- and inter-molecular contacts, in the following CG simulations.

### Principal Component Analysis

Principal component analysis (PCA) on a simulation trajectory of a large biomolecule is used to extract large-scale collective motion of the biomolecule from its small and random internal motion^[38]^. This method comprises the following steps. (1) All conformations in the trajectory are fitted to a reference structure, in order to eliminate the overall translation and rotation of the biomolecule. (2) A covariance matrix of positional fluctuation is constructed, σ_*ij*_ = ⟨ (*r*_*i*_ − ⟨*r*_*i*_⟩)(*r*_*j*_ − ⟨ *r*_*j*_⟩) ⟩, where *r*_*i*_ and *r*_*j*_ represent Cartesian coordinates of the particles selected for PCA, and ⟨⟩ means the coordinate average. (3) The covariance matrix is diagonalized to obtain eigenvectors (PCA modes) and corresponding eigenvalues. Those PCA modes with the largest eigenvalues generally describe collective motion of the biomolecule^[39]^. For each system, 1,000 conformations were contained in the CG trajectory, and the P particles were used to construct the covariance matrix.

To clear see the motion along certain PCA modes, one can project the conformations in the trajectory onto these modes individually. The root mean square inner product (RMSIP) between two sets of PCA modes can measure their dynamic similarity^[40]^:

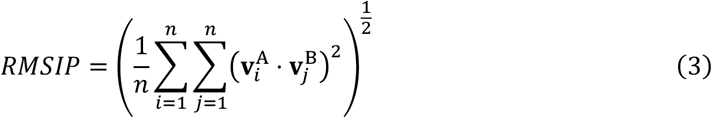

where *v*_*i*_ and *v*_*j*_ represent the *i*^*th*^ and the *j*^*th*^ PCA mode obtained from two different sets of PCA modes (A and B), and *n* is the number of the PCA modes used to compute RMSIP. Since the PCA modes are normalized, an RMSIP of 1 means that the two sets of PCA modes are identical, and an RMSIP of 0 indicates that they are orthogonal.

## Results and Discussion

### Structural dynamics in the four-stage assembly of the 30S subunit

To measure conformational changes of the 16S rRNA in the CG simulations, the root mean squared deviations (RMSD) of all the P particles were calculated, using the native structure of the 16S rRNA in the 30S subunit as the reference. In the CG simulation of the naked 16S rRNA, the system is very unstable with RMSD more than 30.0 Å (**Fig. 2**, black). The tertiary structure of the 16S rRNA is unfolded. Both the platform and the head is expanded, and the junction between the head and the body is extremely mobile (**Fig. 3a**). The results indicate that, without the S proteins, the 16S rRNA itself cannot maintain its native structure.

**Figure 2.**
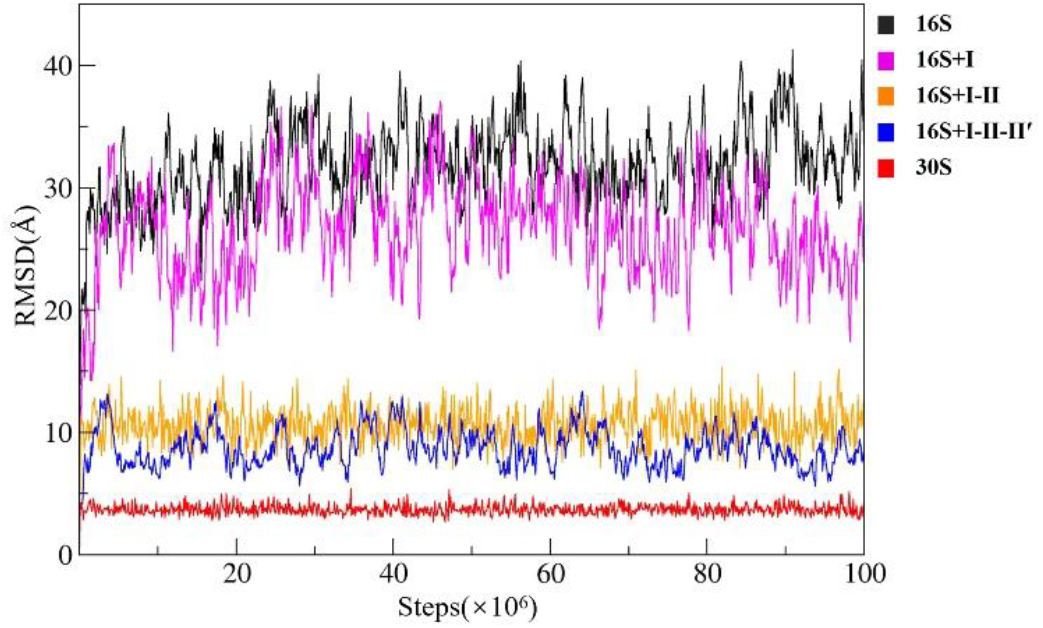
Time evolution of RMSD during the CG simulations. Black: the naked 16S rRNA, magenta: the 16S rRNA with the early-assembly proteins, orange: the 16S rRNA with the early- and mid-assembly proteins, blue: the 16S rRNA with early-, mid-, and mid-late-assembly proteins, and red: the 30S subunit.

**Figure 3.**
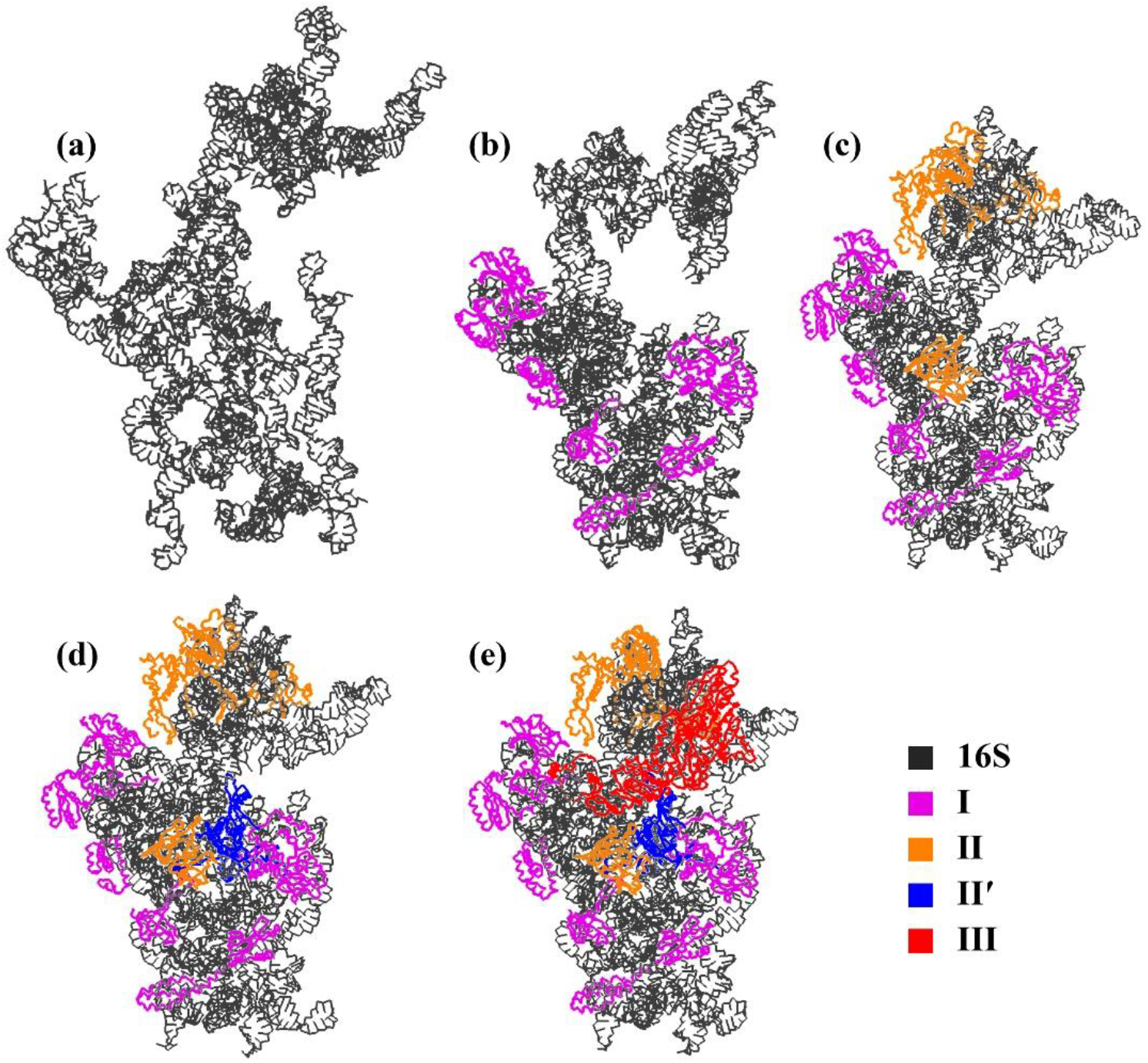
Stability of the 16S rRNA after assembling the S proteins at each stage sequentially. (a) The naked16S rRNA. (b) The 16S rRNA with only the early-assembly proteins. (c) The 16S rRNA with the early- and mid-assembly proteins. (d) The 16S rRNA with the early-, mid- and mid-late-assembly proteins. (e) The 30S subunit. At each stage, the final structure of the CG simulation is shown.

We now want to investigate how the overall stability of the 16S rRNA changes after the S proteins at each stage added sequentially. When only the early-assembly proteins are assembled to the 16S rRNA (denoted as 16S+I), the tertiary structure is still not stable, with the RMSD values essentially between 20-30 Å (**Fig. 2**, magenta). These early-assembly proteins bind to the 5’and the central domain (**Fig. 3b**, magenta), which make up the body and the platform, respectively. Therefore, the body is stabilized and the platform moves close to the body at the early stage compared to the naked 16S rRNA (**Fig. 3a**). However, no early-assembly protein is assembled to the 3’major domain that makes up the head, so the head is nearly as mobile as that in the naked 16S rRNA. This may explain why the RMSD values of the 16S+I system are still large. After the mid-assembly proteins are assembled (denoted as 16S+I-II), the stability of the system is greatly increased. The RMSD values fluctuate around 10.0 Å (**Fig. 2**, orange). According to the positions of the five mid-assembly proteins on the 16S rRNA (**Fig. 3c**, orange), four of them are located on the head that join the two halves of the 3’ major domain together, therefore, the mobility of the head is significantly decreased. Only one mid-assembly protein (S8) is located at the back of the body. The two mid-late assembly proteins are close to the central junction between the body and the head (**Fig. 3d**, blue). Once they are assembled (denoted as 16S+I-II-II’), the stability is increased a little (**Fig. 2**, blue) compared to the 16S+I-II system (**Fig.2**, orange). The 16S+I-II-II’ system is also called 21S reconstitution intermediate^[20]^, which lacks most of the head-body contacts and the beak region in the head is flexible. The late-assembly proteins would make additional connections between the head and the body, and restrict the motion of the beak (**Fig. 3e**, red). Therefore, in the final 30S subunit, the 16S rRNA is well folded with an average RMSD of 3.7 Å (**Fig. 2**, red).

The assembly order of the S proteins are basically from 5’to 3’of the 16S rRNA^[41]^, which however does not mean the early-assembly proteins would contribute most to the stability of the system. It has been found that, the mid- and the late-assembly proteins play the most important roles in the assembly process, due to their locations on the 16S rRNA.

We conducted PCA on the CG trajectory of the 30S subunit, and only P particles were used to construct the covariance matrix. Conformations in the trajectory were then projected onto an 2D subspace defined by the first two PCA modes (denoted as PC1 and PC2). The sampled region of the 30S subunit is centered around (0 Å, 0Å) (**Fig. 4A**, red). For each PCA mode, we took conformations with the most negative and the most positive projection values, superimposed them, and then drew arrows between the corresponding particles. Along the PC1, the movement of the body tail (spur) is significant (**Fig. 4b**), whereas along the PC2, the open/close movement between the head and the body is pronounced (**Fig. 4c**).

**Figure 4.**
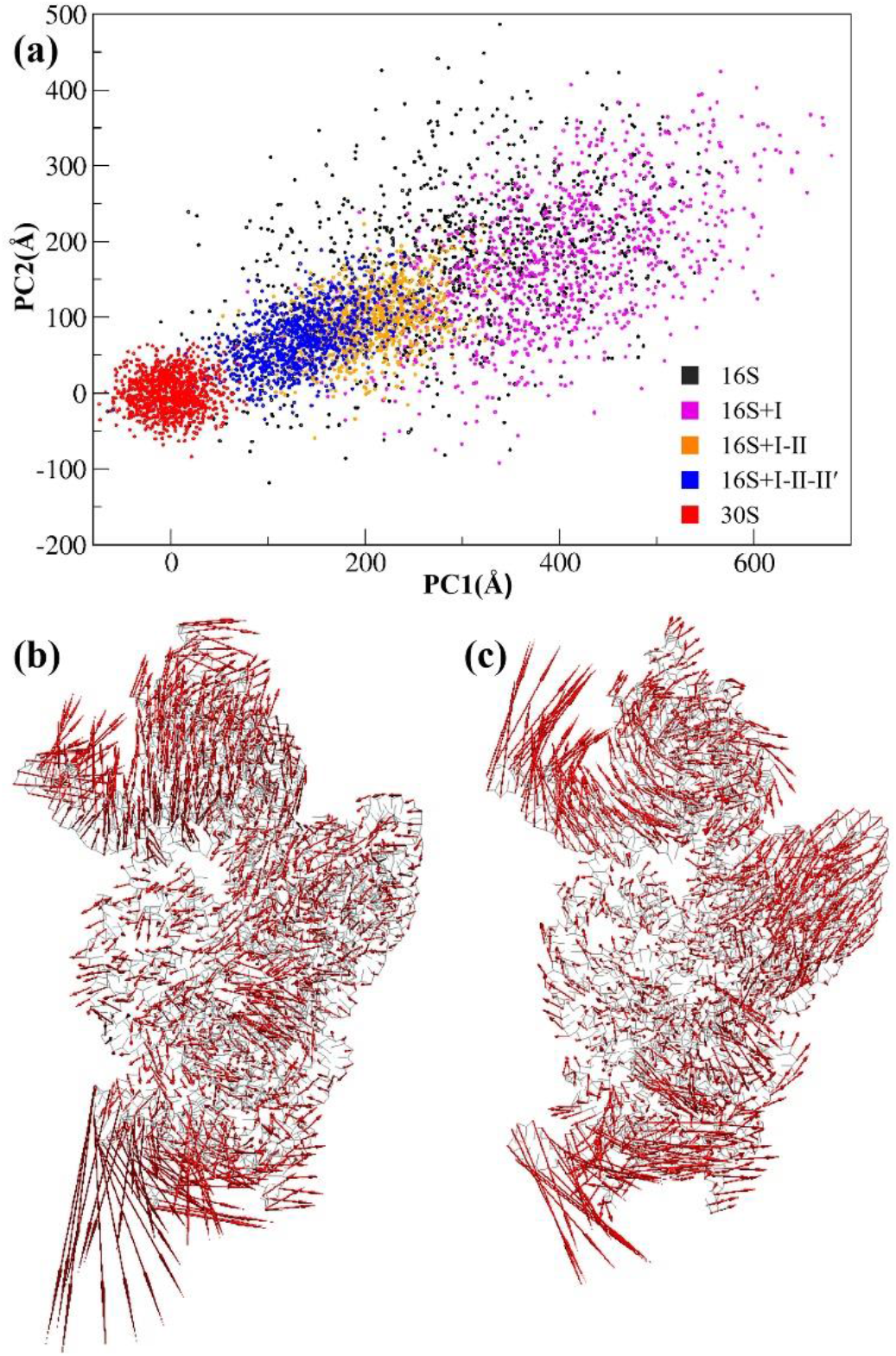
An assembly path of the 30S subunit shown on a subspace defined by the PCA modes. PCA was conducted on the CG trajectory of the 30S subunit, and only P particles were used to construct the covariance matrix. The first two PCA modes with the largest eigenvalues (PC1 and PC2) were chosen to define the 2D subspace. (a) Projections of the different CG trajectories onto the subspace. (b) Collective motion along the PC1. (c) Collective motion along the PC2.

The other CG trajectories were also projected onto the same subspace. The naked 16S rRNA wildly samples a very large conformational space (**Fig. 4a**, black). With the S proteins at the different stages assembled, one can clearly see an ‘assembly path’ on the subspace. At the early stage, although the sampled space is still quite large (**Fig. 4a**, magenta), it is somewhat narrower than that of the naked 16S rRNA (**Fig. 4a**, black). At the mid-stage, the conformational space is sharply decreased (**Fig. 4a**, orange), and at the mid-late-stage (**Fig. 4a**, blue), the conformational space is getting close to that of the folded 16S rRNA (**Fig. 4a**, red).

### The role of specific S proteins in the assembly

After investigating the S proteins at the four stages, another question is that, what is the specific role of individual S protein in the assembly of the 30S subunit? We have carried out a series of CG simulations starting from the naked 16S rRNA, then adding the S proteins one by one following the order shown in Table 1, and finally ending to the 30S subunit. From S4 to S20 that are the early-assembly proteins, the systems are all very mobile with the RMSD values fluctuating around 30.0 Å (**Fig. 5a**, dashed curves in magenta). When S7 and S8, the first two added mid-assembly proteins, are assembled, the RMSD values are only decreased a little (**Fig. 5a**, the two dashed curves in orange essentially between 20-25 Å). Once S9 is assembled, the system is significantly stabilized with the RMSD values suddenly dropping to around 10.0 Å (the solid curve in orange). S9 has a long C-terminal tail inserting deeply into the head **(Fig. 5b**), which can join the two halves of the 3’ major domain together and reduce the mobility of the head significantly. The assembly of the two mid-late-assembly proteins (S5 and S12) does not decrease the RMSD much (**Fig. 5a**, dashed curves in blue). When S2, the first added late-assembly protein, is assembled, the RMSD values are decreased to about 7.0 Å (**Fig. 5a**, the dashed curve in red). However, once S3 is assembled, the RMSD values again have a drop to about 4.0 Å (**Fig. 5a**, the solid curve in red). S3 spans the gap between the beak in the head and the shoulder in the body (**Fig. 5c**), so it can stabilize the conformation of the beak and make contacts between the head and the body. The assembly of last three late-stage proteins (S10, S14, and S21) do not lower the RMSD values anymore (**Fig. 5a**, dashed curves in red). The separated RMSD values suggest key events in the assembly process of the 30S subunit. The first event is the folding of the head starting with the assembly of S9, and the second event is the restricted domain motion between the head (beak) and the body starting with the assembly of S3. That is to say, the two dividing S proteins (S9 and S3) play important roles in the assembly of the 30S subunit, due to their unique locations on the 16S rRNA (**Fig. 5b-5c**).

**Figure 5.**
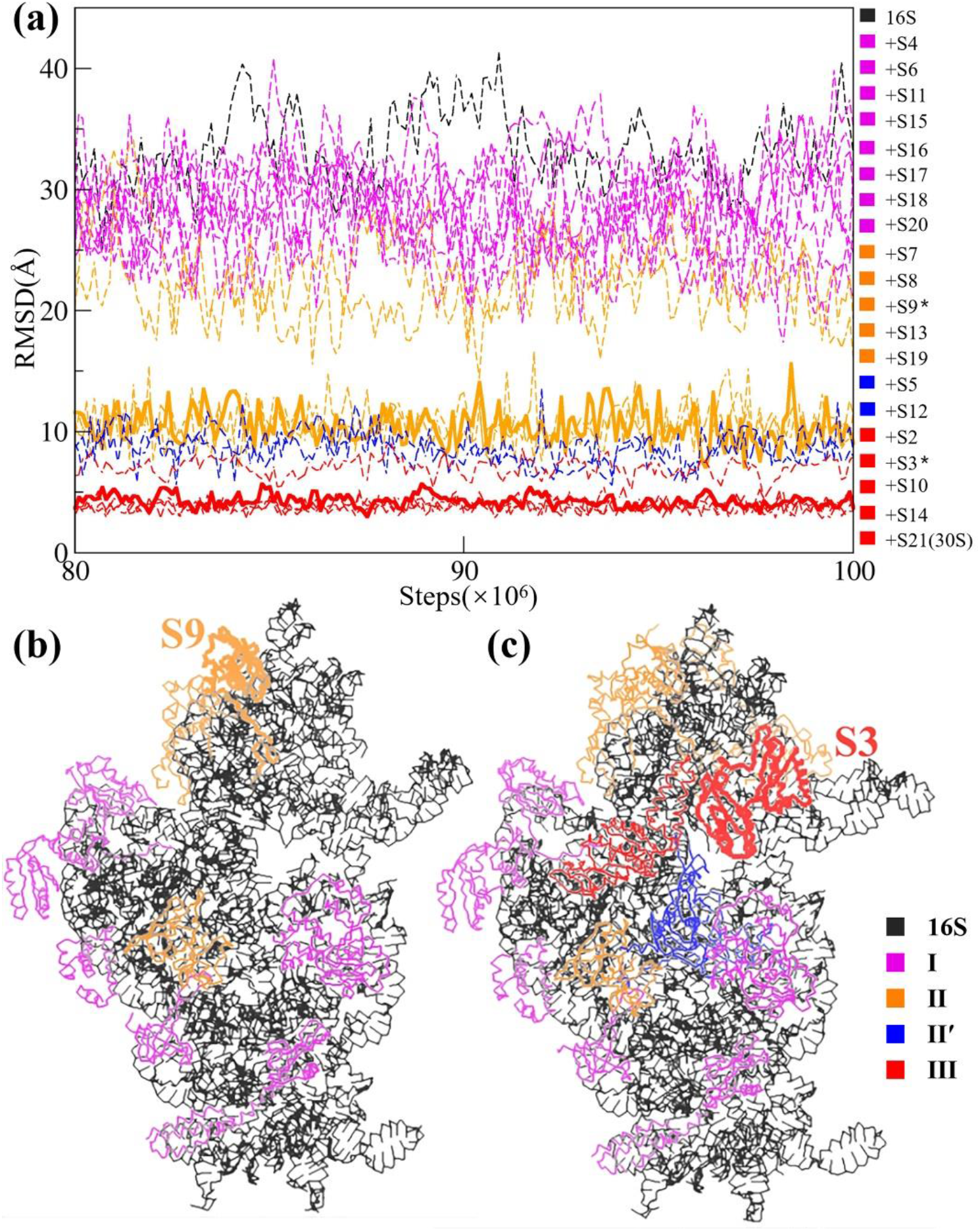
The role of individual S protein in the assembly of the 30S subunit. (a) Equilibriumed (from 80-to 100-million steps of the CG simulations) RMSD values of all the simulated systems. Starting from the naked 16S rRNA, then adding the S proteins one by one following the order shown in Table 1, and finally ending to the 30S subunit. The RMSD curves at the four stages are colored differently. In the color bar at the right side, the two dividing proteins (S9 and S3) are labeled by *. The two corresponding RMSD curves are shown by solid lines while other curves are shown by dashed lines. (b) The position of S9. (c) The position of S3.

To quantify how the 16S rRNA obtains its collective motion in the 30S subunit during the assembly process, PCA was performed on every CG trajectory using the P particles to construct the covariance matrix. For each simulated system, the first ten PCA modes with the largest eigenvalues were selected as a set. We used the PCA modes from the 30S subunit as a reference set, and then computed RMSIP (Eq. 3) between any other set of the PCA modes and the reference (**Fig. 6**). The RMSIP between the PCA modes of the naked 16S rRNA and the reference is only 0.48, indicating that the conformational dynamics of the naked 16S rRNA is very different to that in the 30S subunit. The early-assembly proteins and the first two added mid-assembly proteins do not increase RMSIP much, which keeps around 0.55. Once S9 is assembled, the RMSIP is significantly increase to more than 0.8. The assembly of S9 would make the head essentially fold (**Fig. 5b**), and thus the collective motion of the head relative to the body becomes dominant as that in the 30S subunit. The two mid-late-assembly proteins (S5 and S12) do not increase RMSIP much. After S3 is assembled, the RMSIP is getting to 0.9, which means that the collective motion of the system is already very similar to that in the 30S subunit. The results here also support the key events during the assembly process, as shown in **Fig. 5a**.

**Figure 6.**
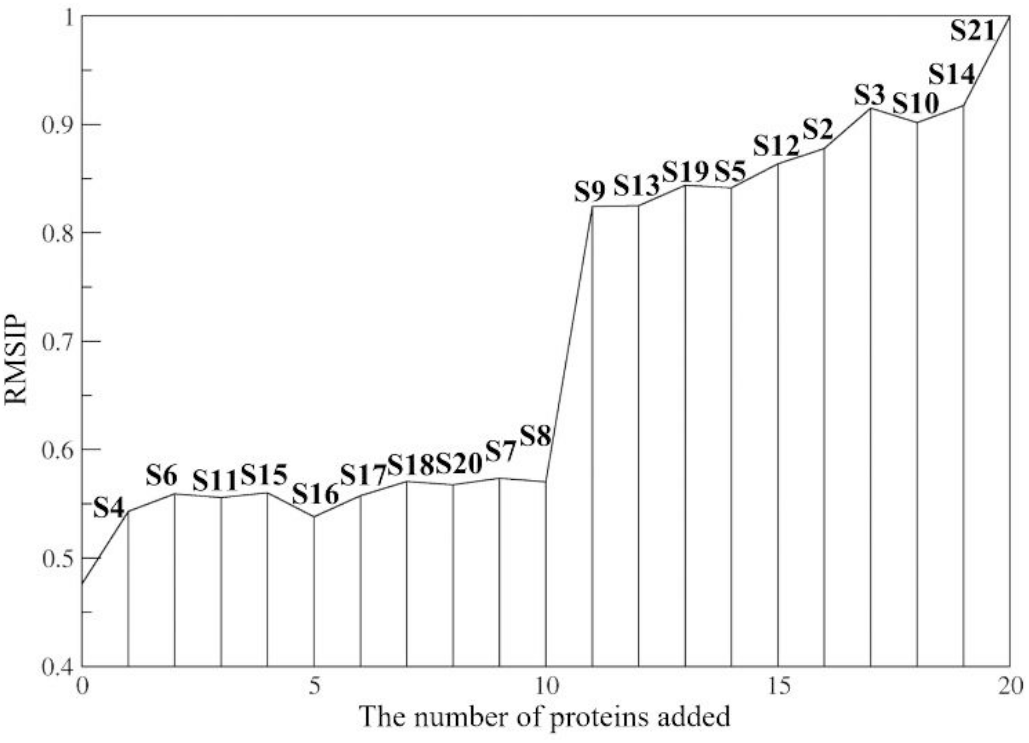
Evolution of the RMSIP in the assembly process of the 30S subunit. The x-axis represents the assembly order of the S proteins shown in Table 1. For each system, the RMSIP was calculated between the PCA modes of the system and those of the 30S subunit. The RMSIP is ending to 1.0 because the system is the 30S subunit after S21 is assembled.

### Structural dynamics in a switched assembly order

What would happen if the assembly order of the S proteins is switched? To address this issue, we have carried out another set of CG simulations by not following the kinetic map (Table 1) but assembling the S proteins in a descending order of their molecular weights (S2, S3, …, S21).

According to RMSD curves shown in Figure 7a, two dividing S proteins, S5 and S11, were detected in this switched assembly order. Once S5 is assembled, the RMSD values significantly drop from around 30.0 Å to around 15.0 Å (**Fig. 7a**, the solid curve in blue). S5 is located at the junction (**Fig. 7b**), which can restrict the motion between the head and the body. However, since there are only four proteins (S2 to S5) assembled at this time, the body and the head are not well folded yet, and the platform is still detached from the body (**Fig. 7b**). When S11 is assembled, the RMSD values are decreased from more than 10.0 Å to about 8.0 Å (**Fig. 7a**, the solid curve in magenta). S11 is binding on the platform, which can significantly stabilize the latter. At this time, S12 to S21 are not assembled yet, which make the body still partially unfolded (**Fig. 7c**). The results indicate that, structural dynamics in the switched assembly order is different to that following the kinetic assembly map (**Fig. 5**). It should be noted that, without knowing the kinetic assembly map beforehand, the number of possible assembly orders is tremendous, and the CG simulations alone have a limited predictive power on determining which assembly order is reasonable. That is to say, the current work can only study structural dynamics during the assembly process of the 30S subunit based on its kinetic assembly map.

**Figure 7.**
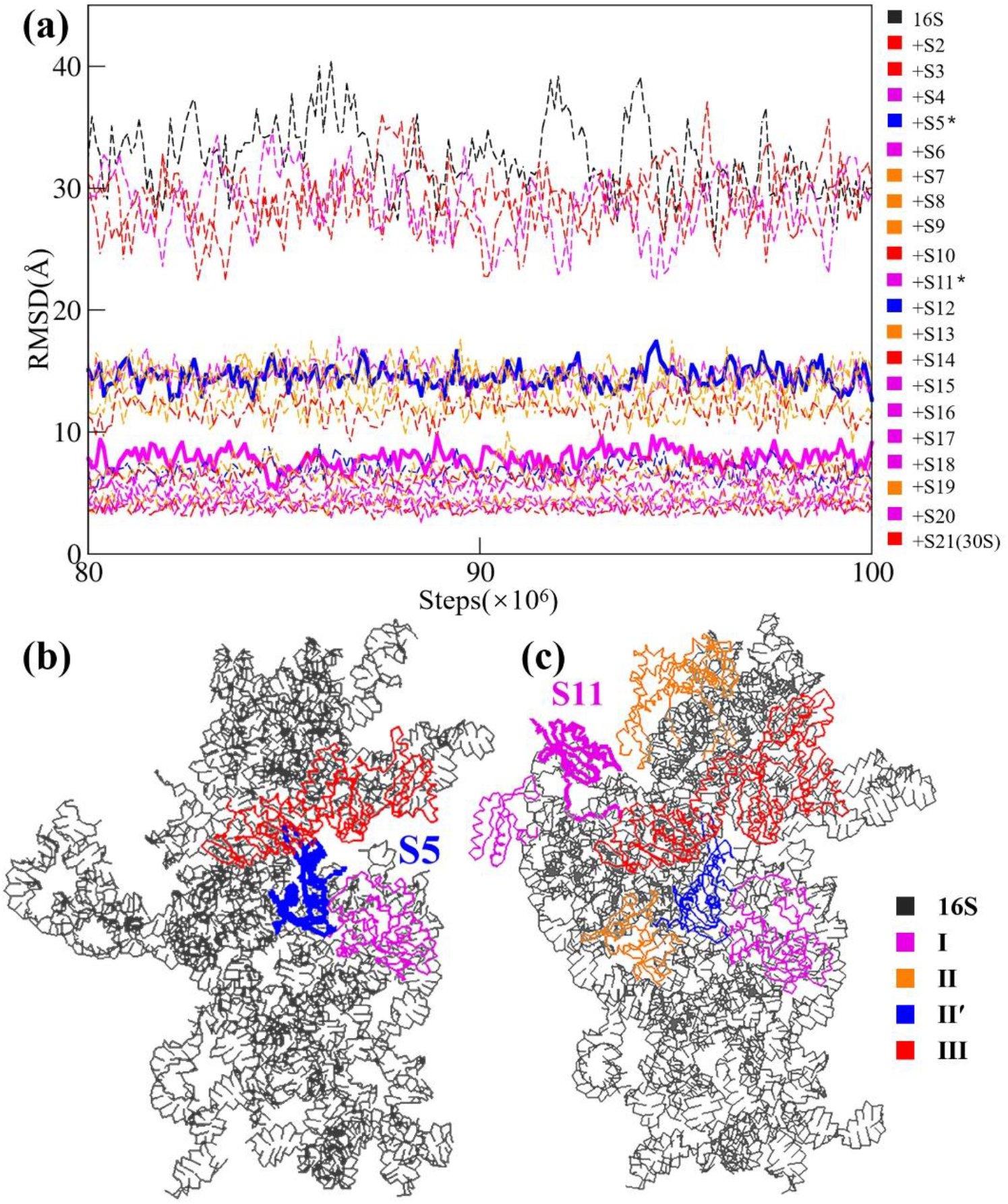
The role of individual S protein in a switched assembly order of the 30S subunit. (a) Equilibriumed (from 80-to 100-million steps of the CG simulations) RMSD values of all the simulated systems. Starting from the naked 16S rRNA, then adding the S proteins one by one in a descending order of their molecular weights, and finally ending to the 30S subunit. The RMSD curves are still colored according to the stages the corresponding S proteins belong to, as in Figure 5a. In the color bar at the right side, the two dividing proteins (S5 and S11) are labeled by *. The two corresponding RMSD curves are shown by solid lines while other curves are shown by dashed lines. (b) The position of S5. (c) The position of S11.

## Conclusions

More and more structures of large biomolecular complexes have been solved, with the development of structural biology techniques, such as cryo-EM. However, with only one static structure, the dynamic process of how different components assemble into a complex is still unclear. As a complement to experiments, computer simulation may serve as an important tool for studying the assembly process of large biomolecular complexes. Due to the generally large size of a biomolecular complex and long time scale of its assembly, CG simulation is an appropriate choice.

This paper works on the 30S small subunit of the *E. coli* ribosome, which is a model system for studying the assembly of biomolecular complexes. Based on the proposed kinetic assembly map, a series of CG simulations have been carried out, from a naked 16S rRNA to a fully assembled 30S subunit. The CG simulations reveal conformational changes at the different assembly stages, and particularly those S proteins with important contributions to the assembly are detected. Our study has proved the rationality of the kinetic assembly map, and in the meantime, provided structural details during the assembly of the 30S subunit.

Our strategy can be applied for studying structural dynamics in the assembly of any large biomolecular complex as long as the assembly map is available. However, due to the high complexity, assembly orders for many biomolecular complexes are unknown. In this case, one may get such information from some advanced experimental techniques like time-resolved cryo-EM^[42]^, or use additional computational tools to predict the assembly order of biomolecular complexes^[43]^.

## Acknowledgements

This work is supported by National Key Research and Development Program of China (2021YFA1301504), National Natural Science Foundation of China (91953101), and Chinese Academy of Sciences Strategic Priority Research Program (XDB37040202). The Supercomputing Center of USTC provides partial computer resources for this project, and we are grateful to Mr. Yundong Zhang for his technical supports.

## Conflict of interest

The authors declare that they have no conflict of interest.

## Notes

### Competing Interest Statement

The authors have declared no competing interest.

